## Supplementary material for "Photodamage repair pathways contribute to the accurate maintenance of the DNA methylome landscape upon UV exposure": Supp Figures and Tables

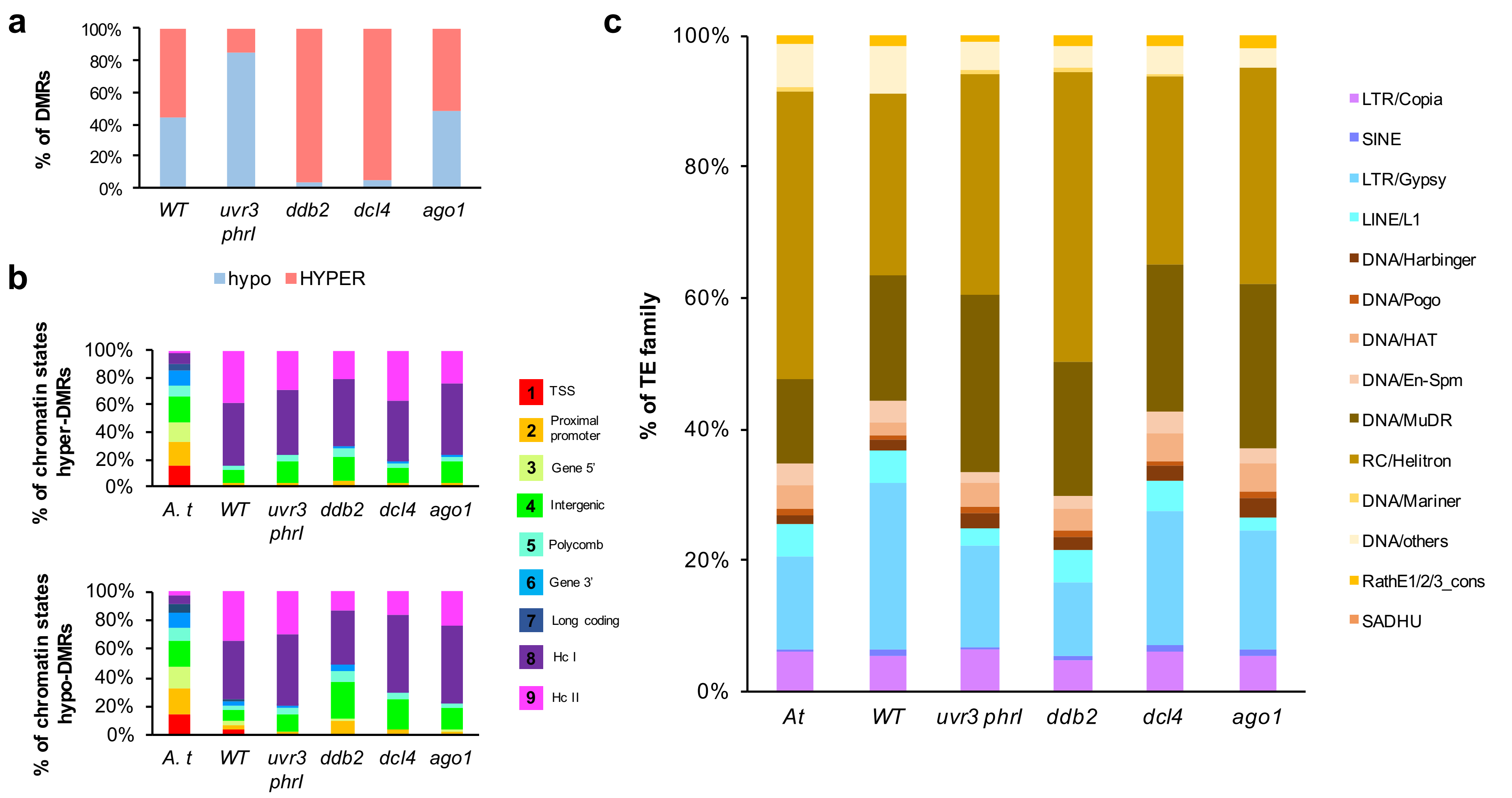


**Supplemental Figure 1**

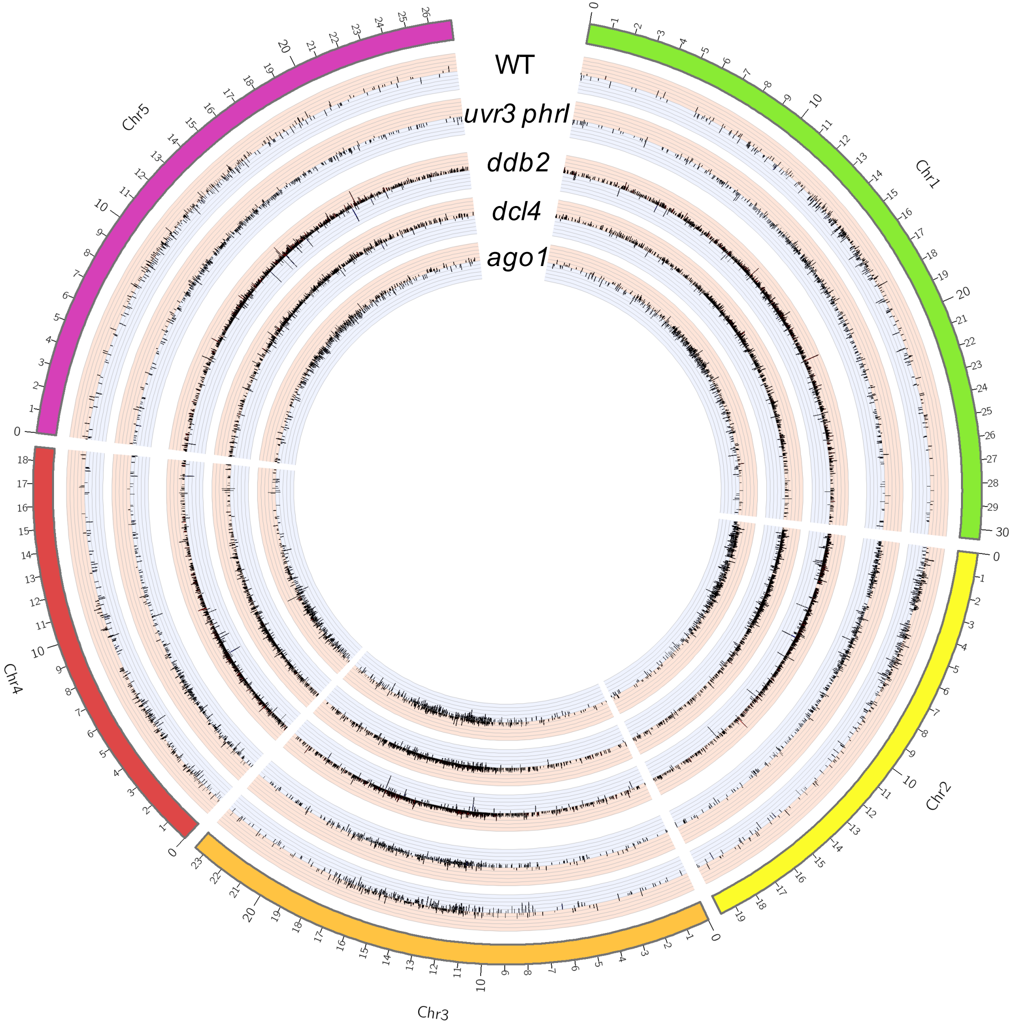


**Supplemental Figure 2**

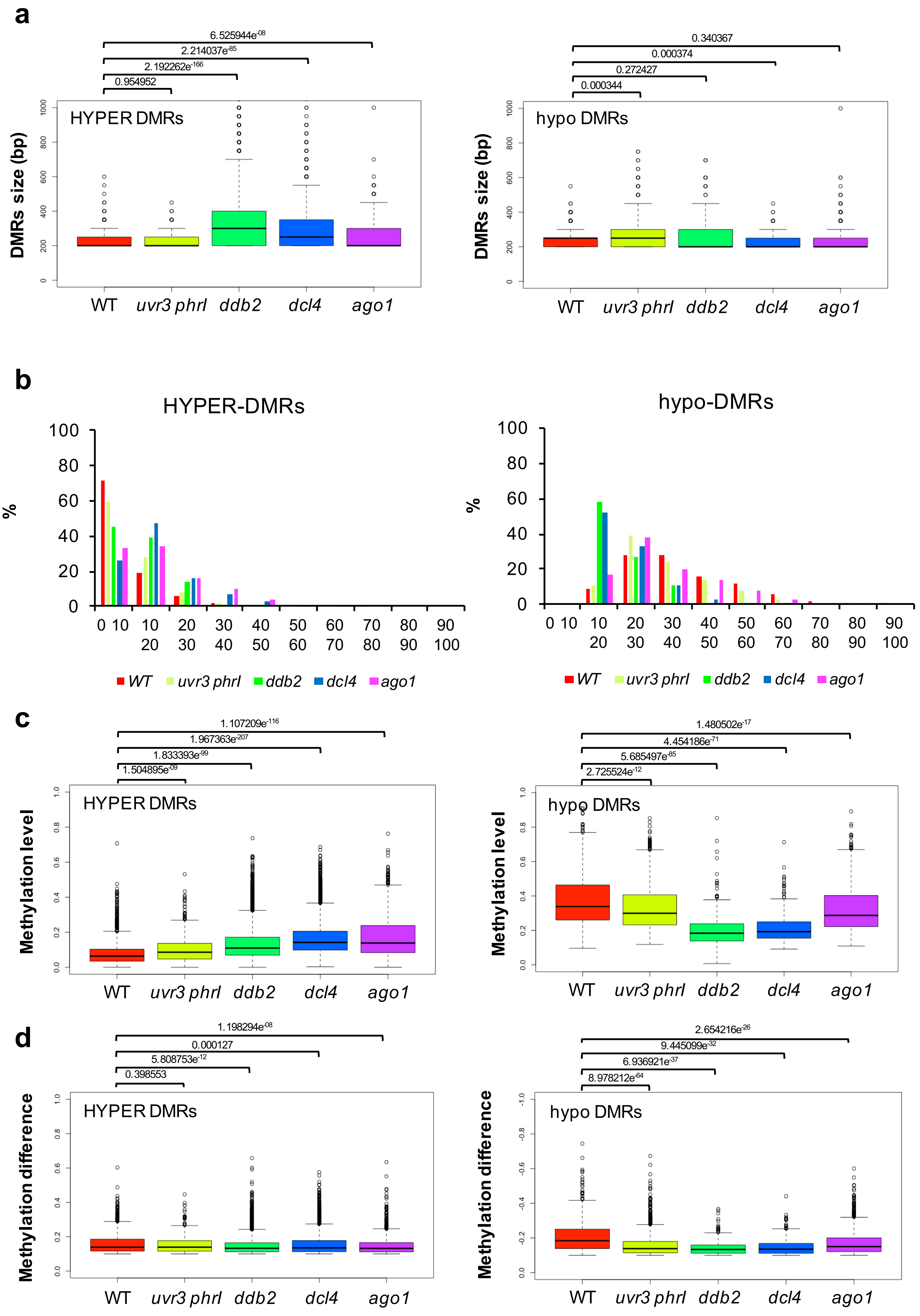


**Supplemental Figure 3**

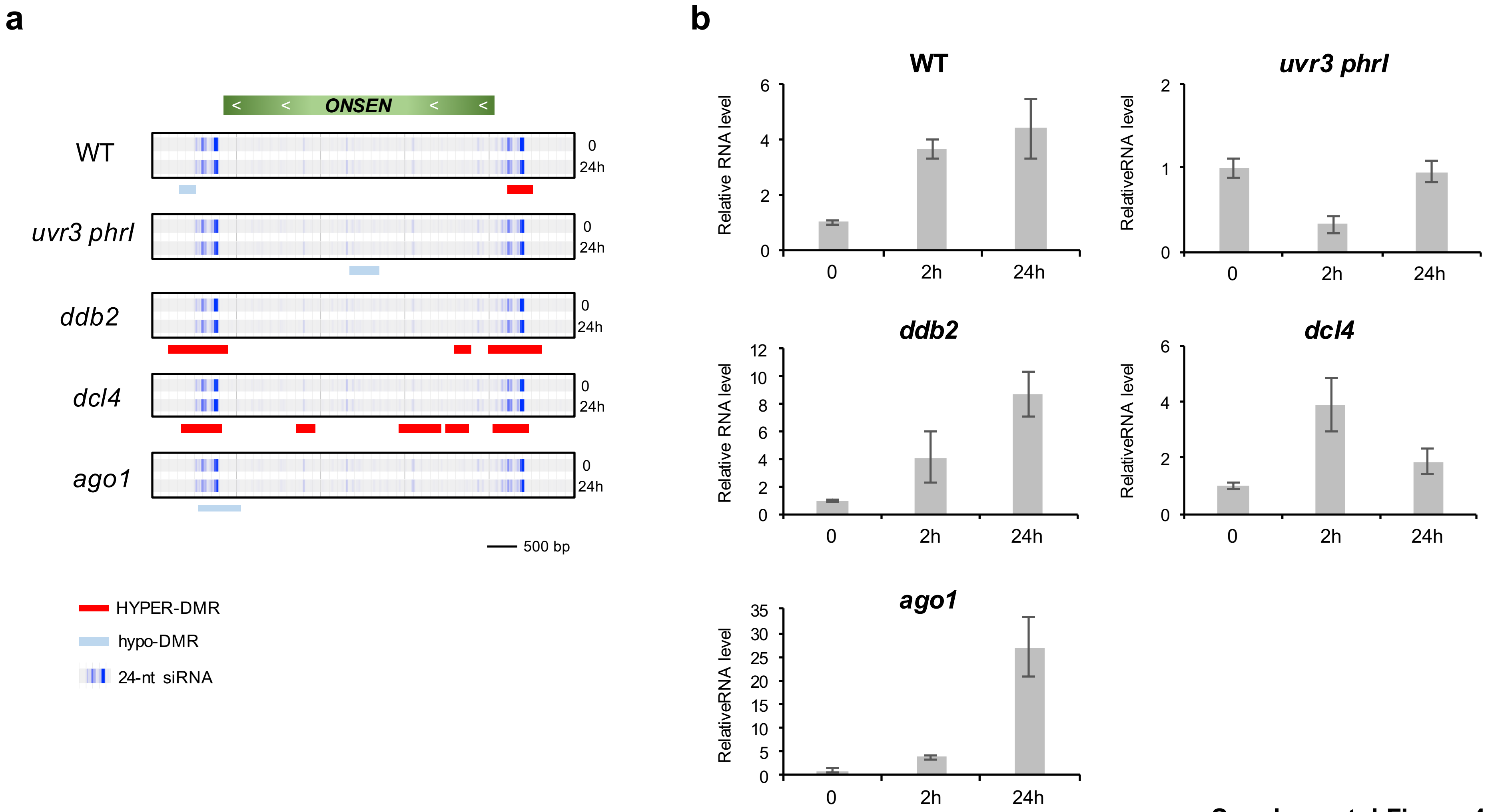


**Supplemental Figure 4**

### HYPER-DMRs

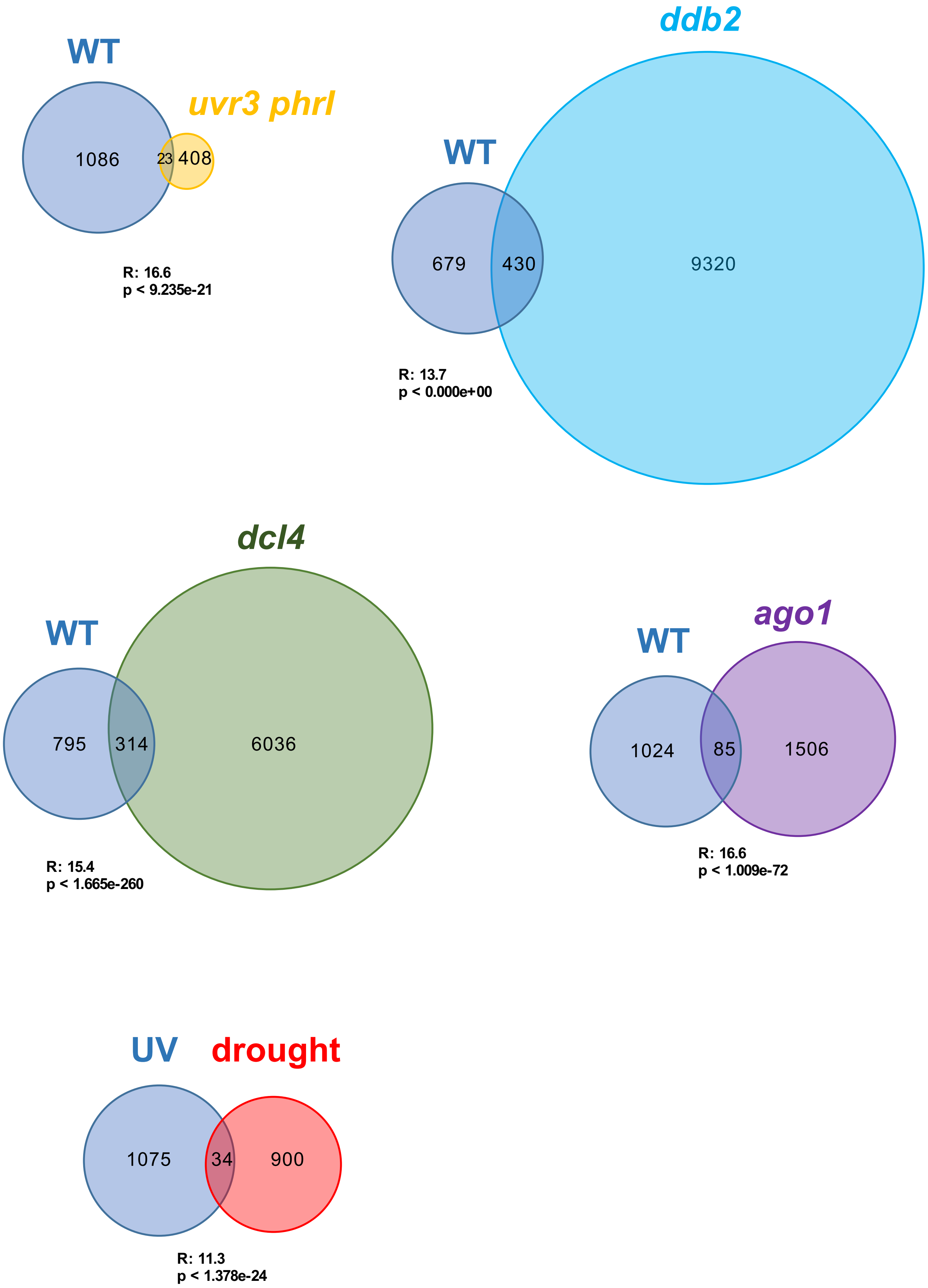


Supplemental Figure 5

#### hypo-DMRs

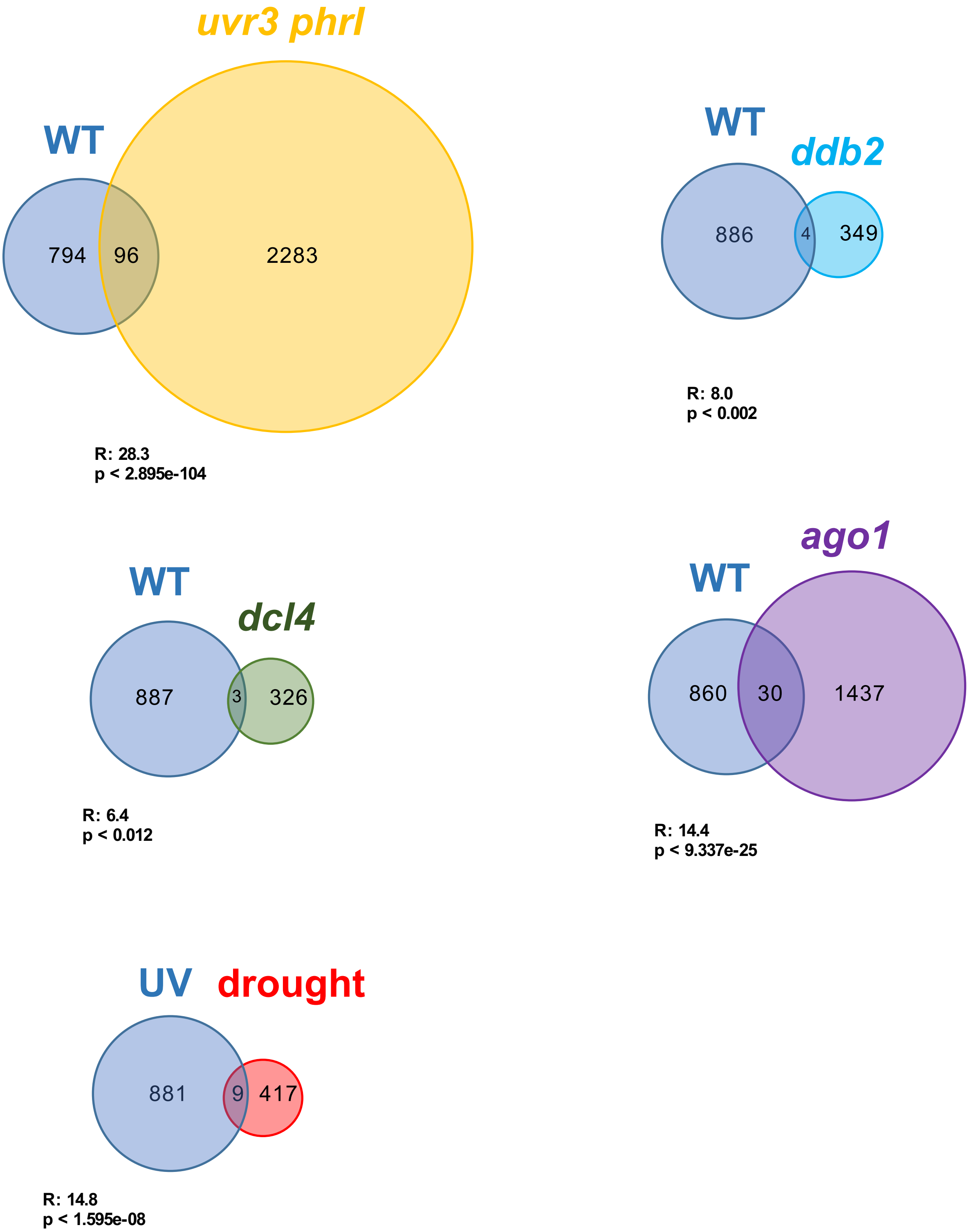


**Supplemental Figure 6**

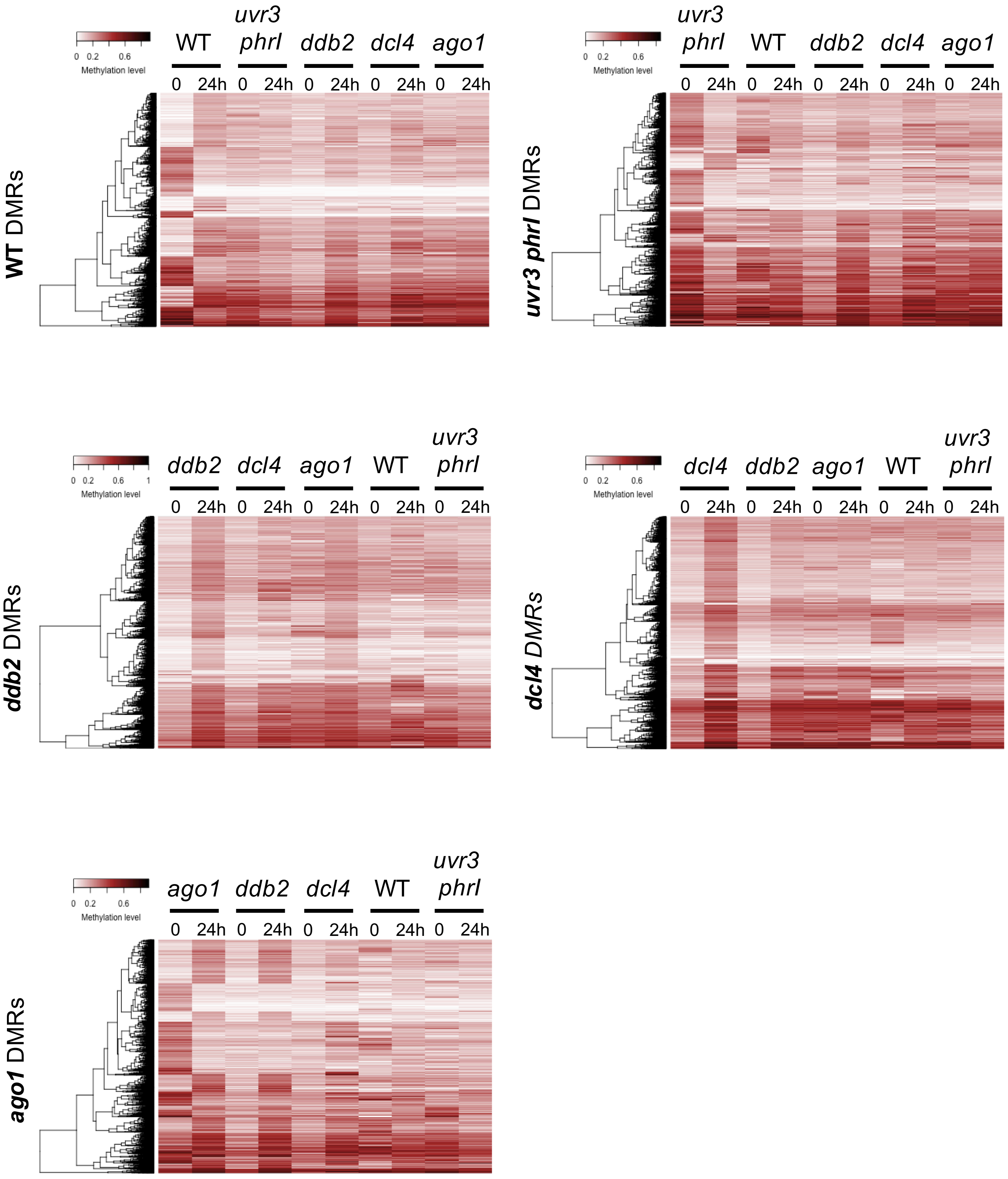


**Supplemental Figure 7**

**Supplemental Figure 8**

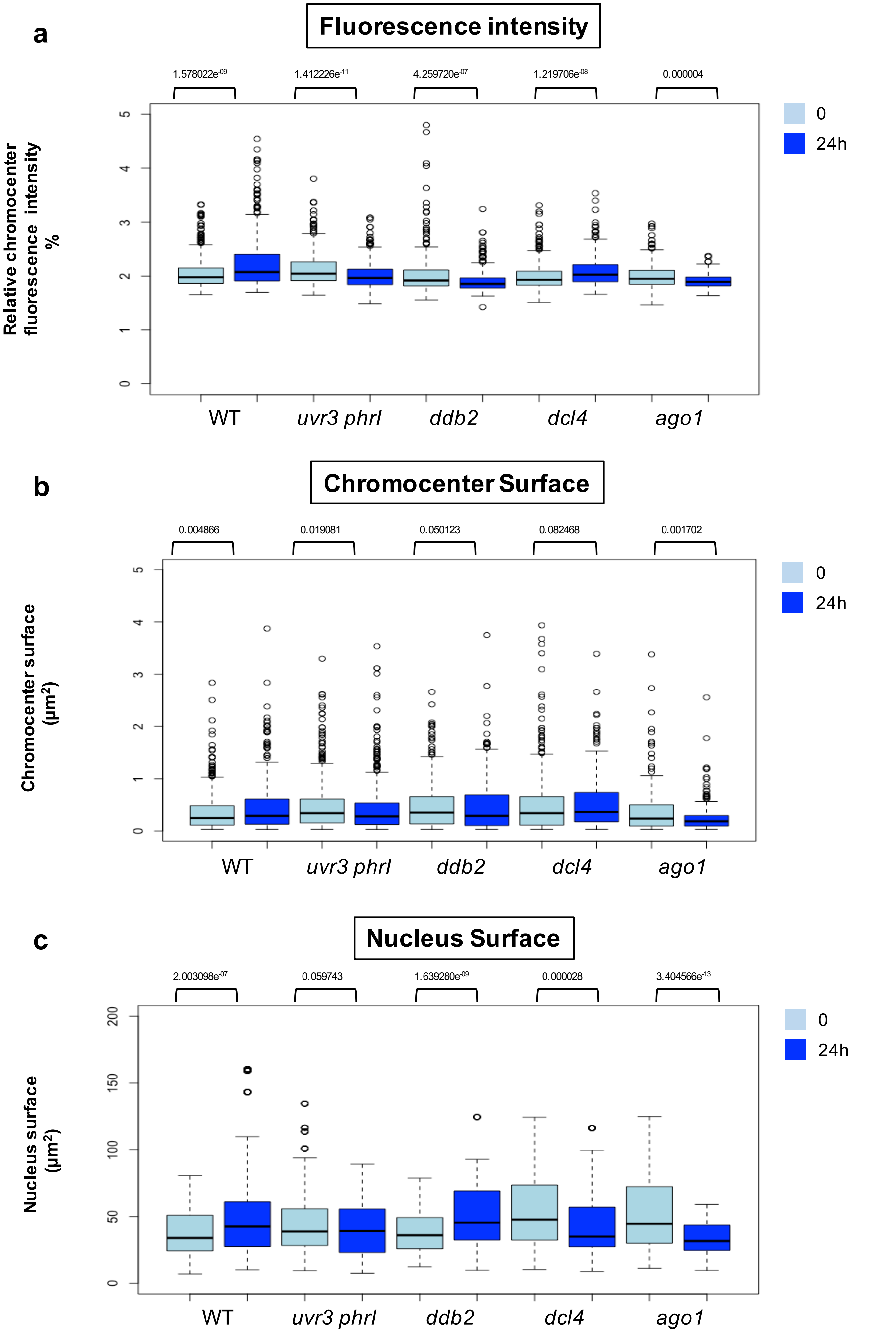

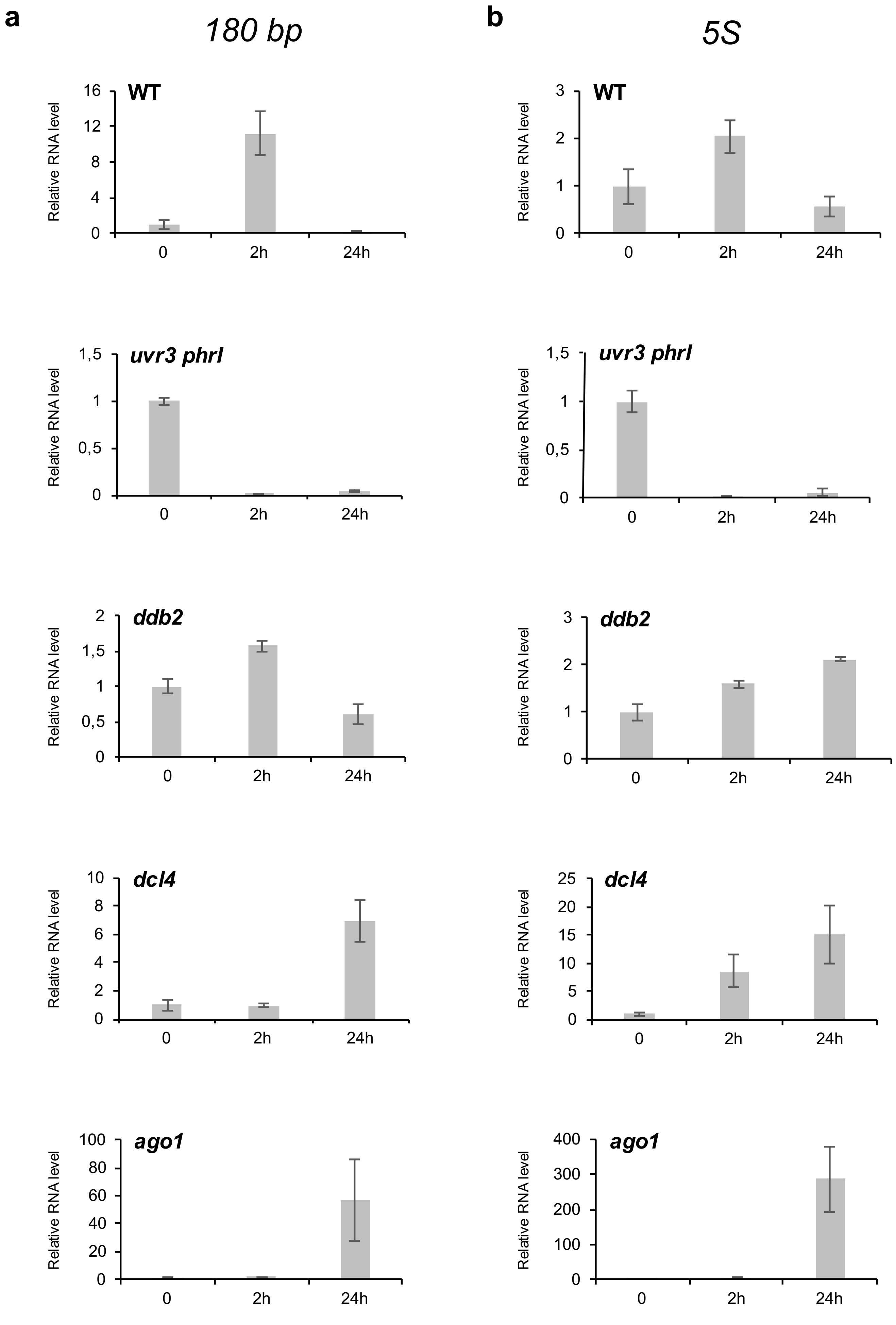


Supplemental Figure 9

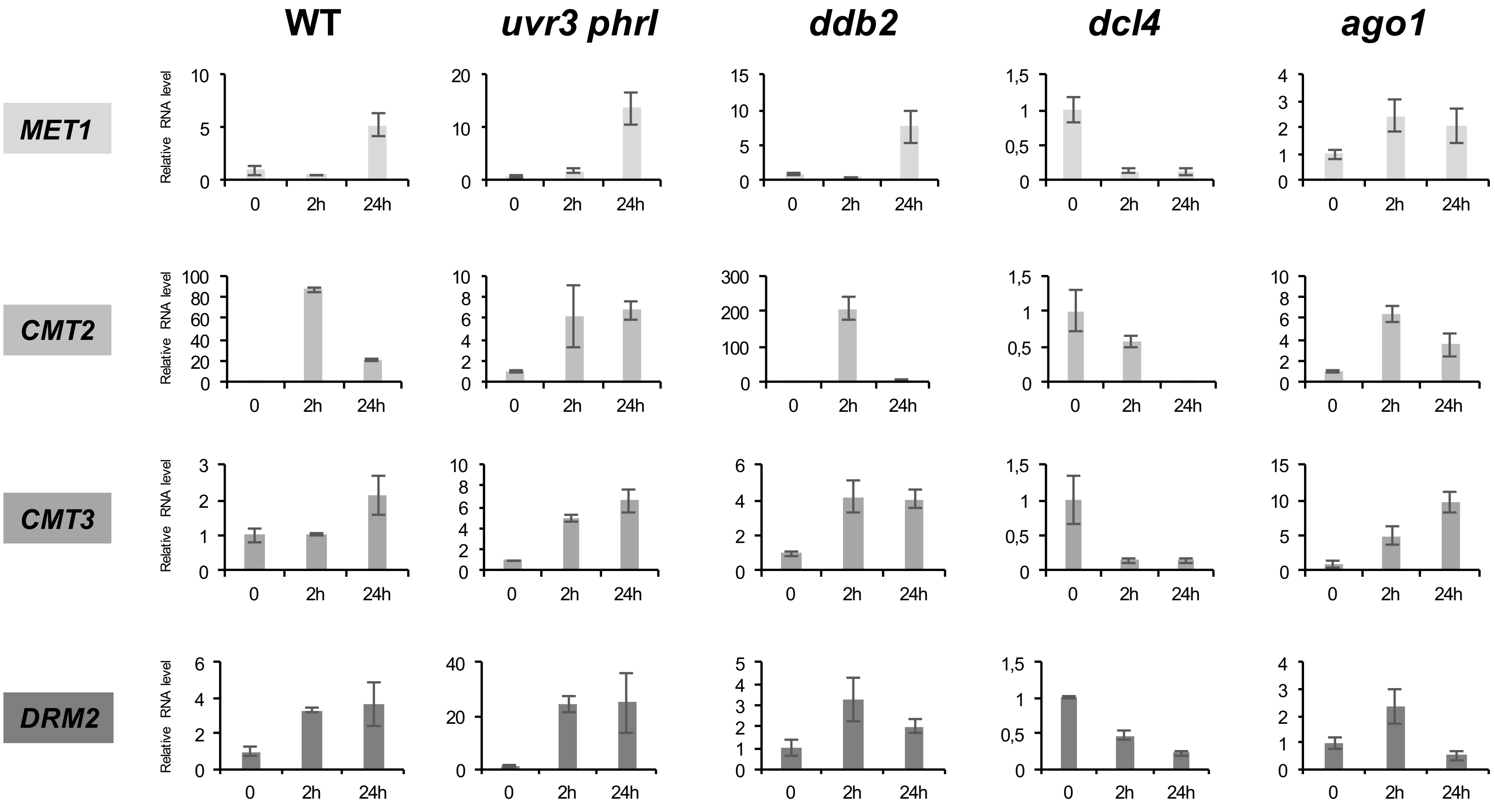


**Supplemental Figure 10**

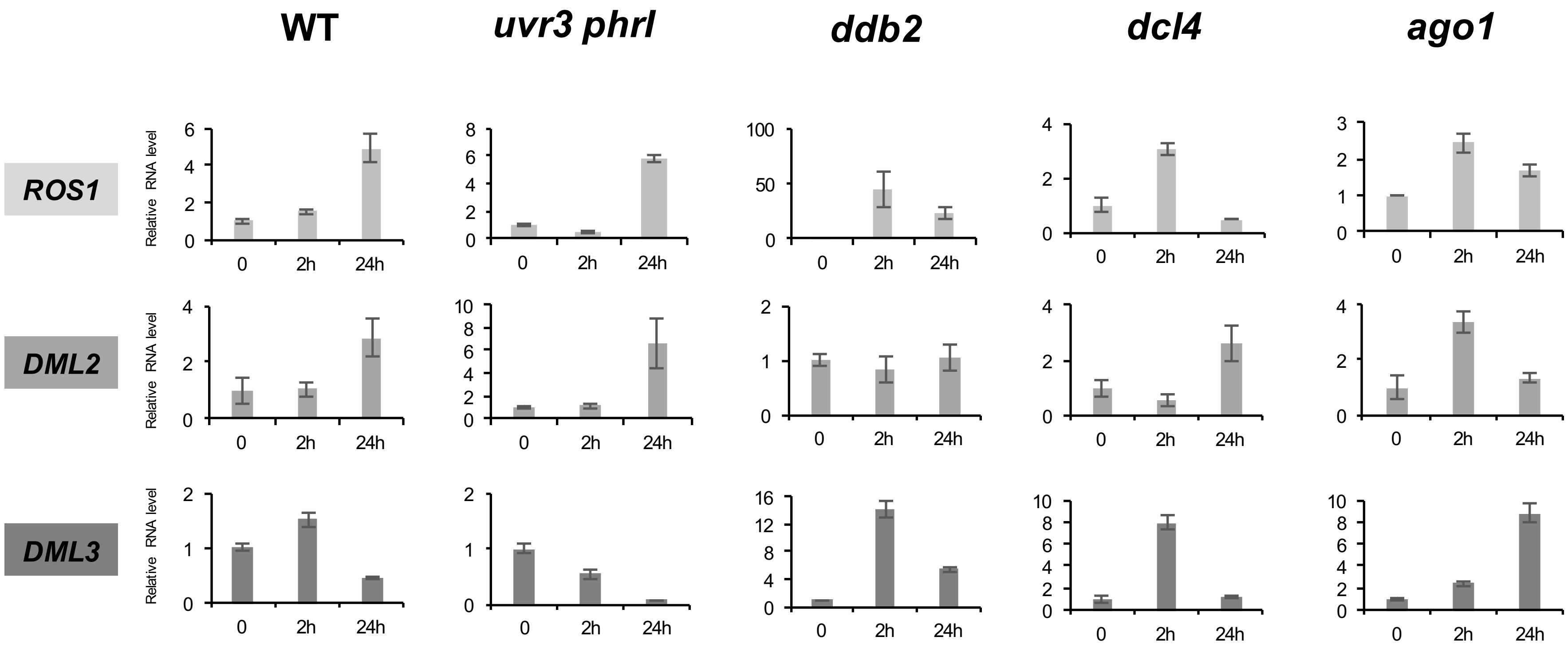


**Supplemental Figure 11**

Supplemental Figure 12

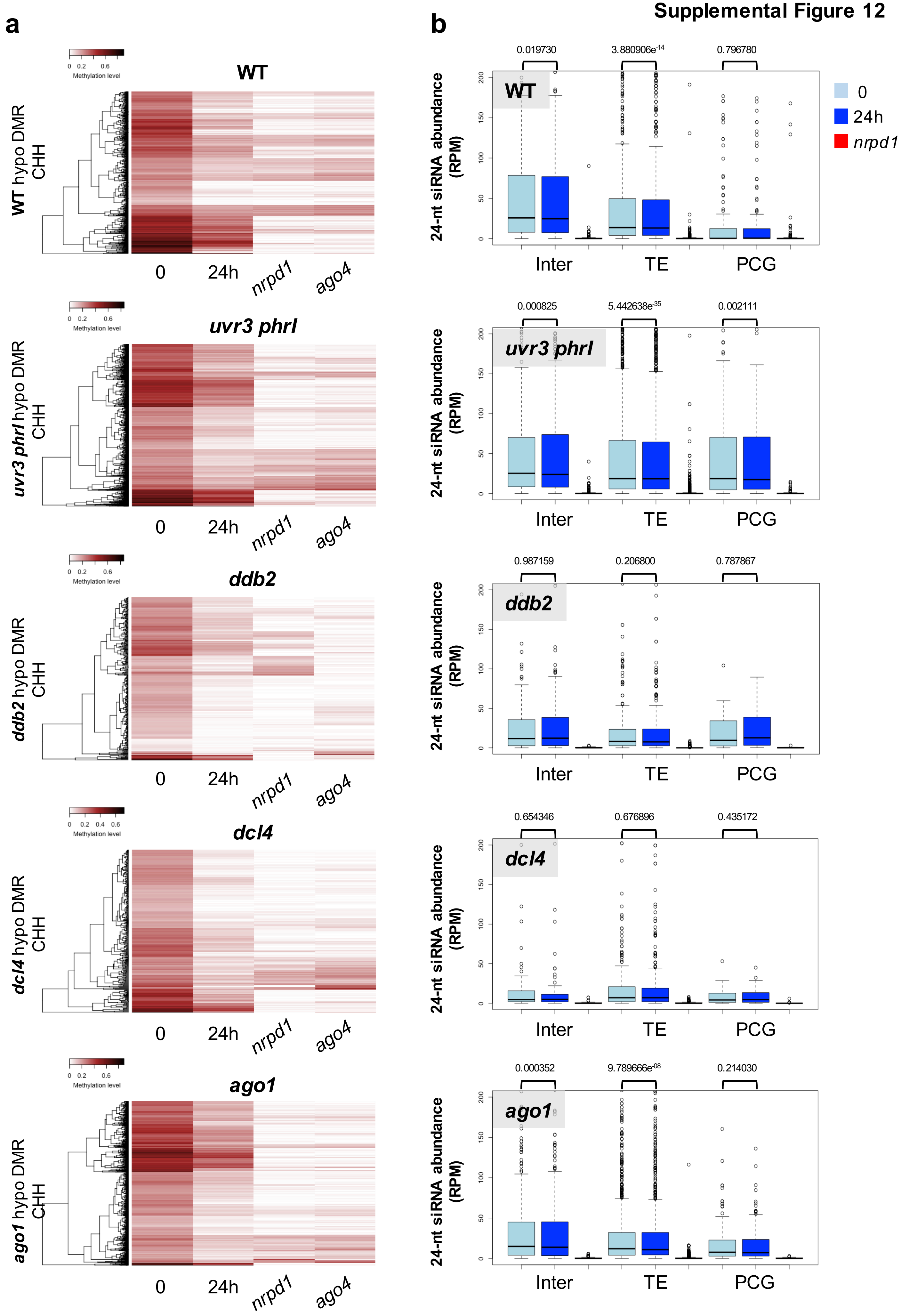

a

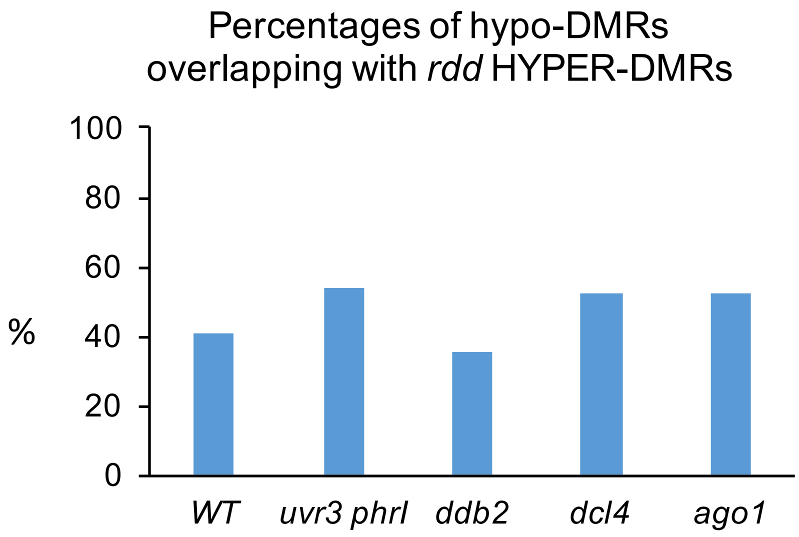


b

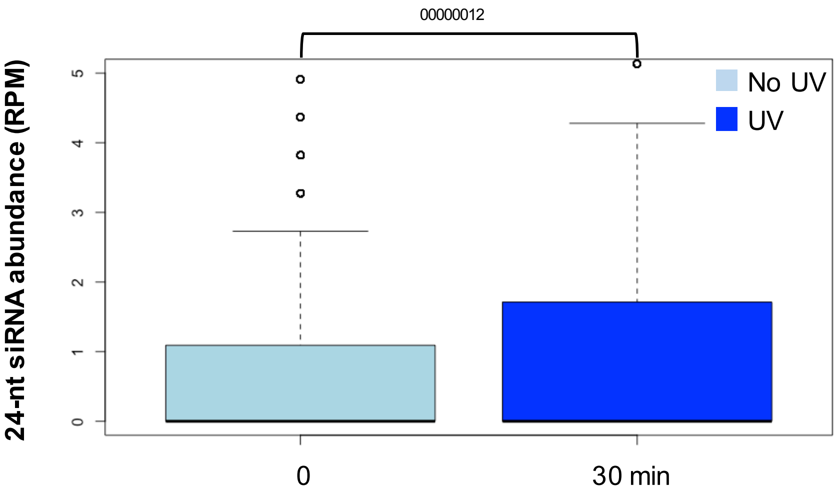


**Supplemental Figure 13**

### Supplemental Figure 14

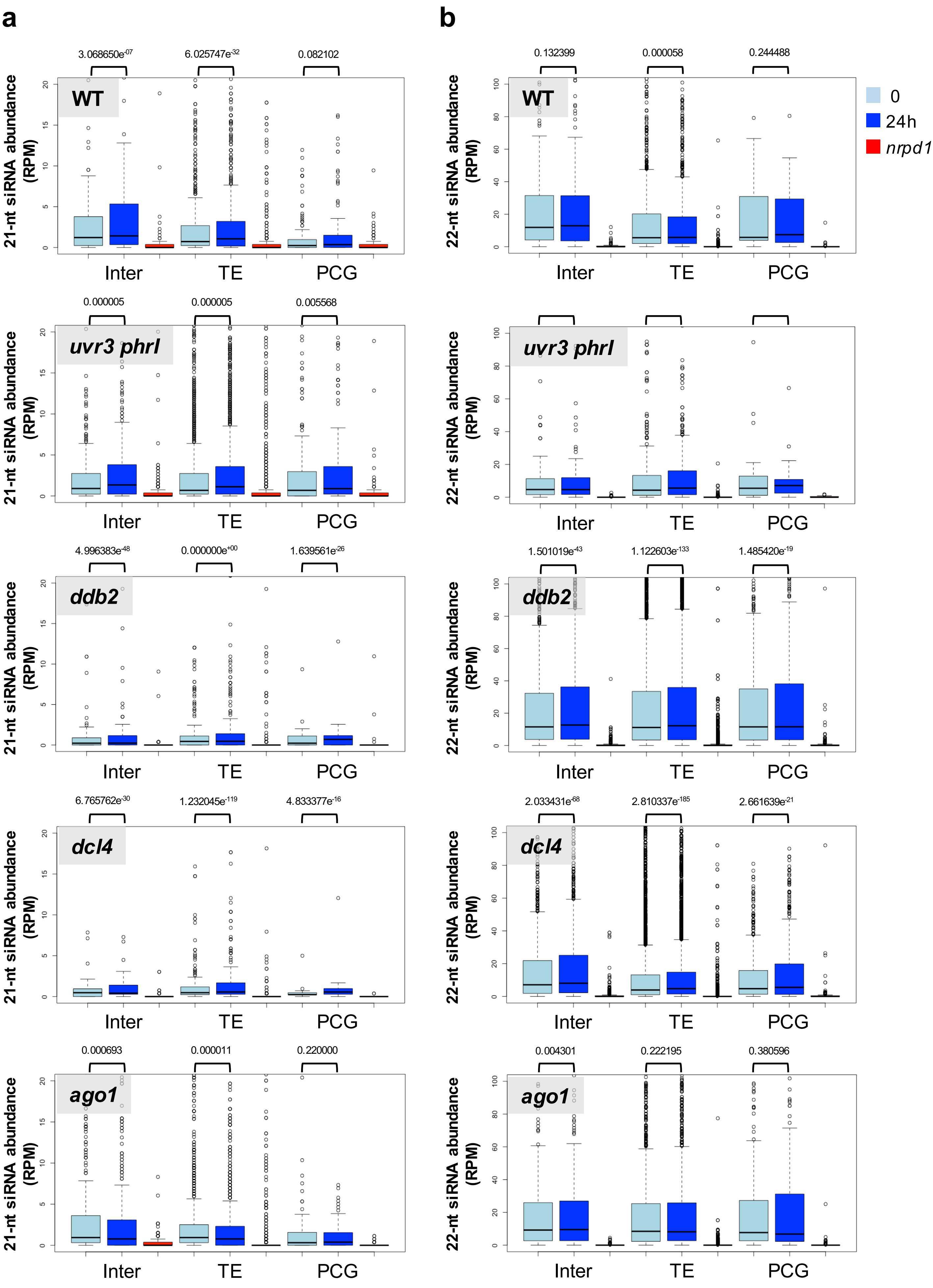

### Supplemental Figure 15

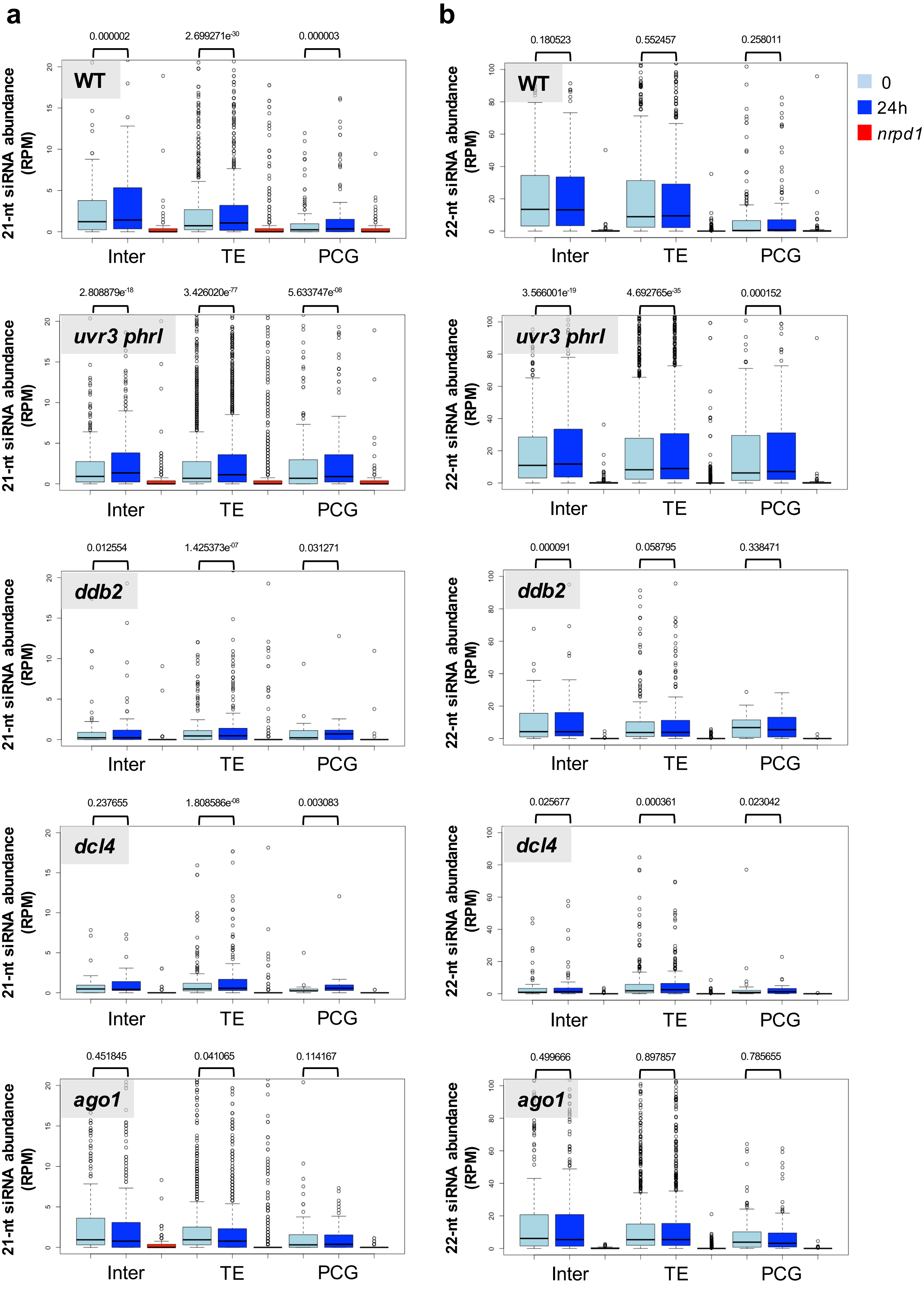

**Supplemental Figure 16**

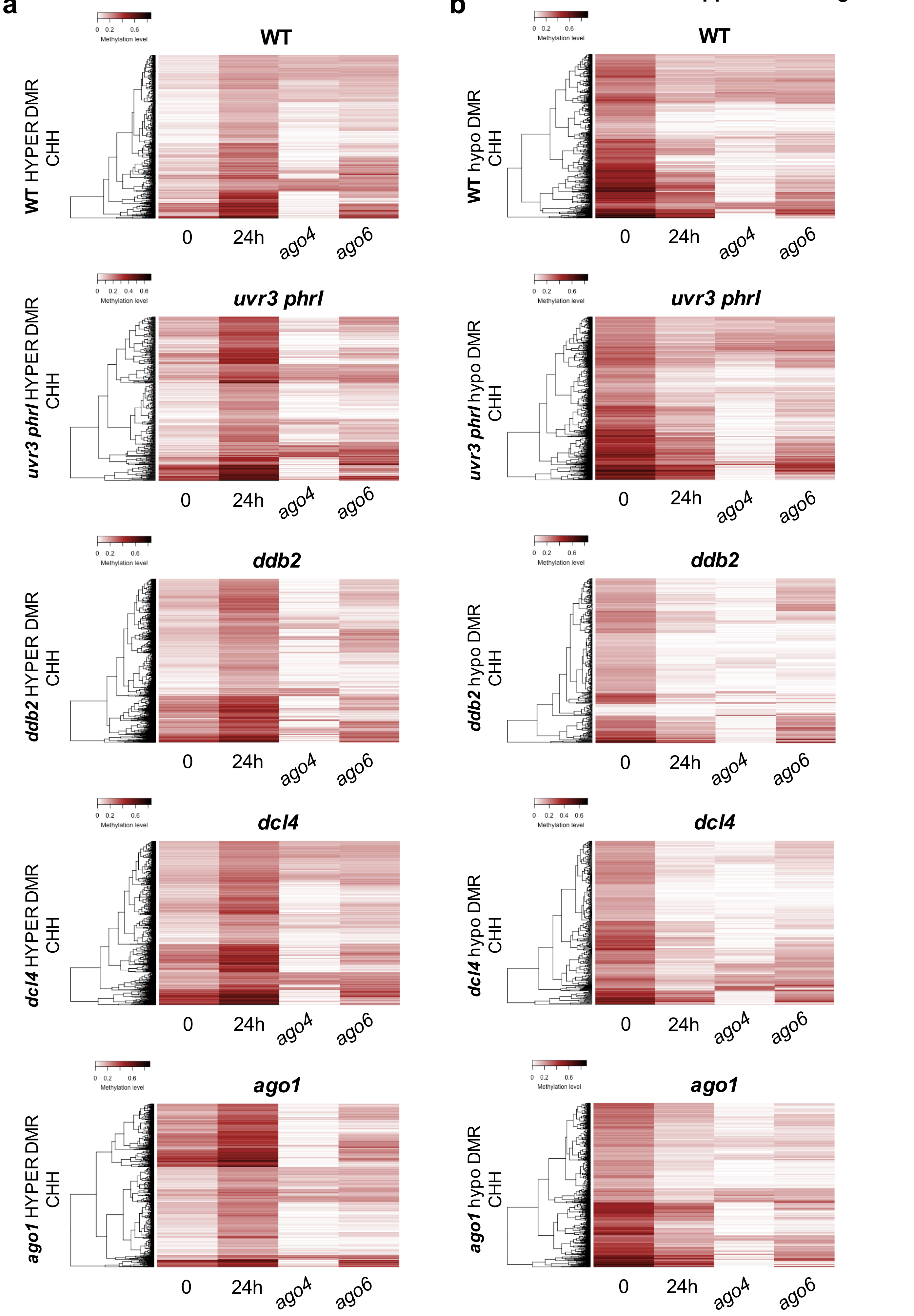

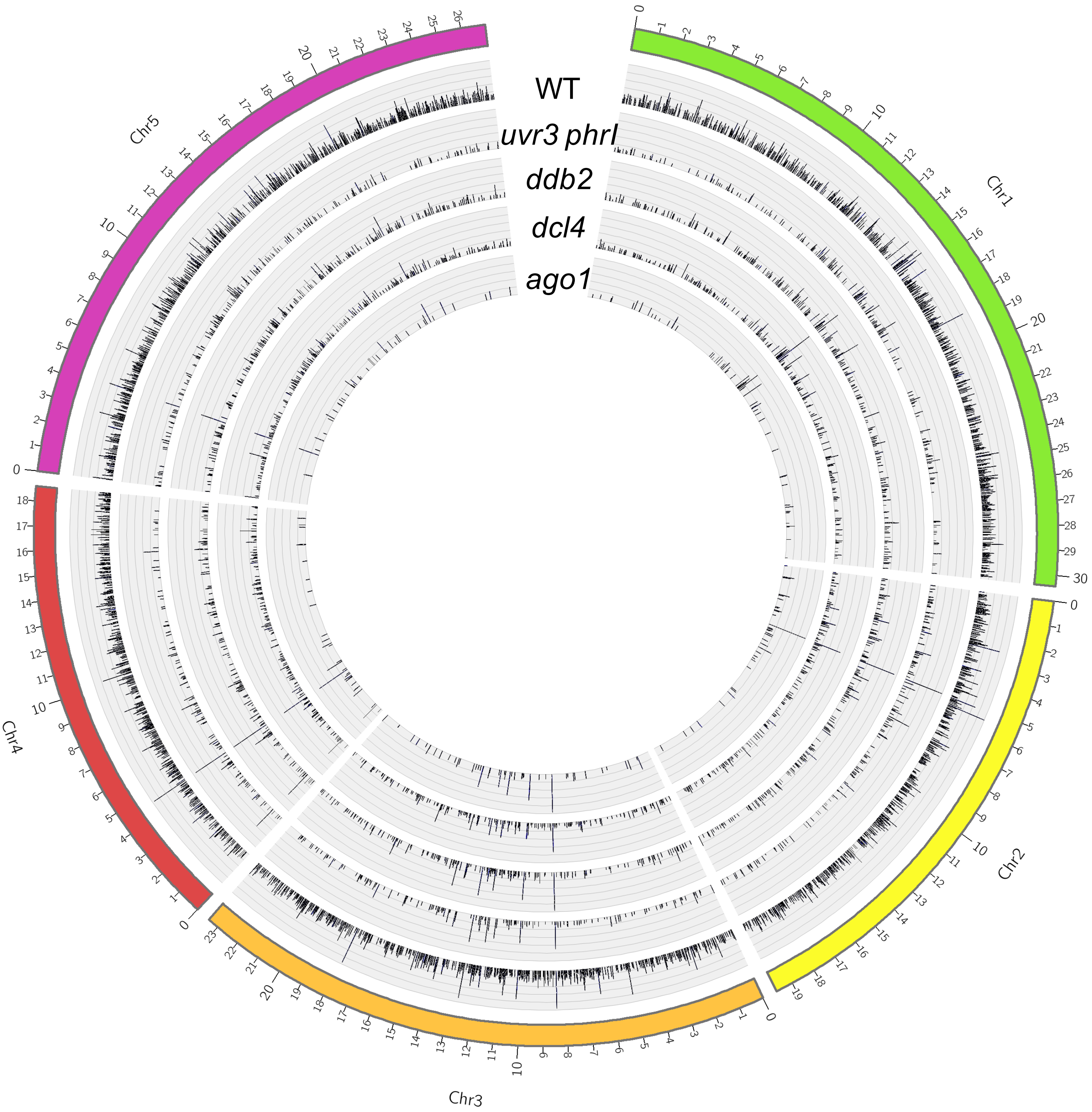


**Supplemental Figure 17**

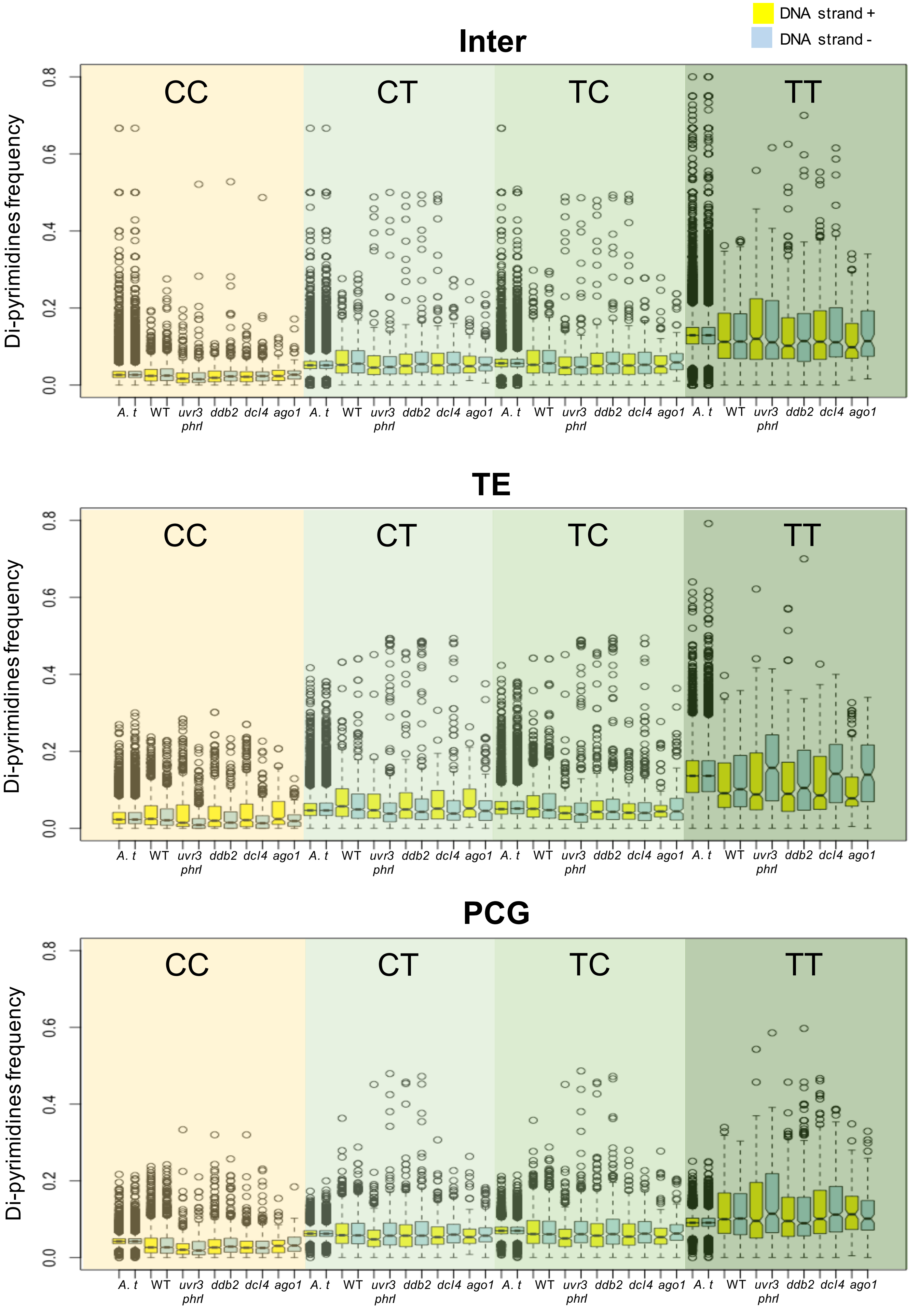


**Supplemental Figure 18**

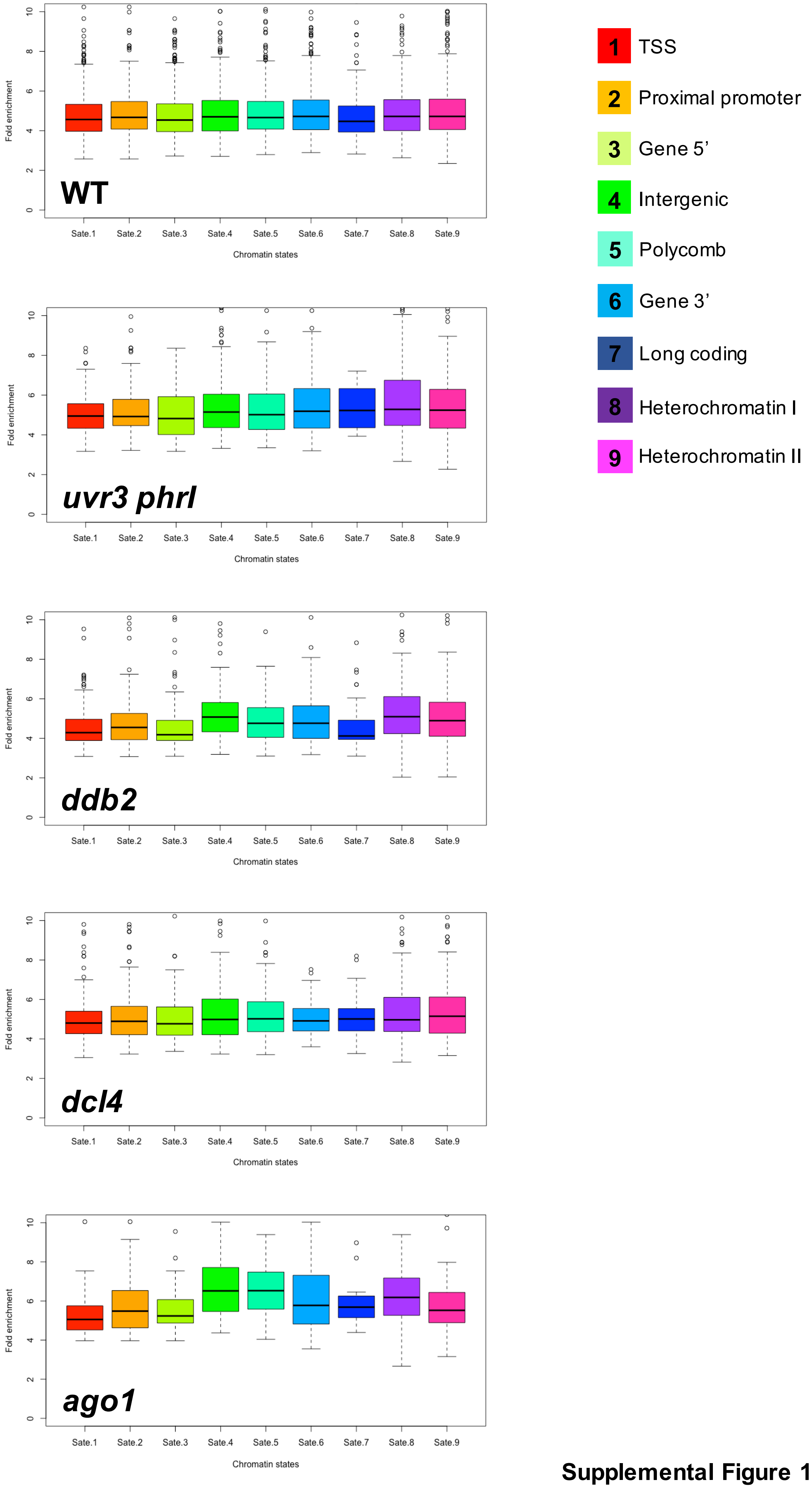


**Supplemental Figure 19**

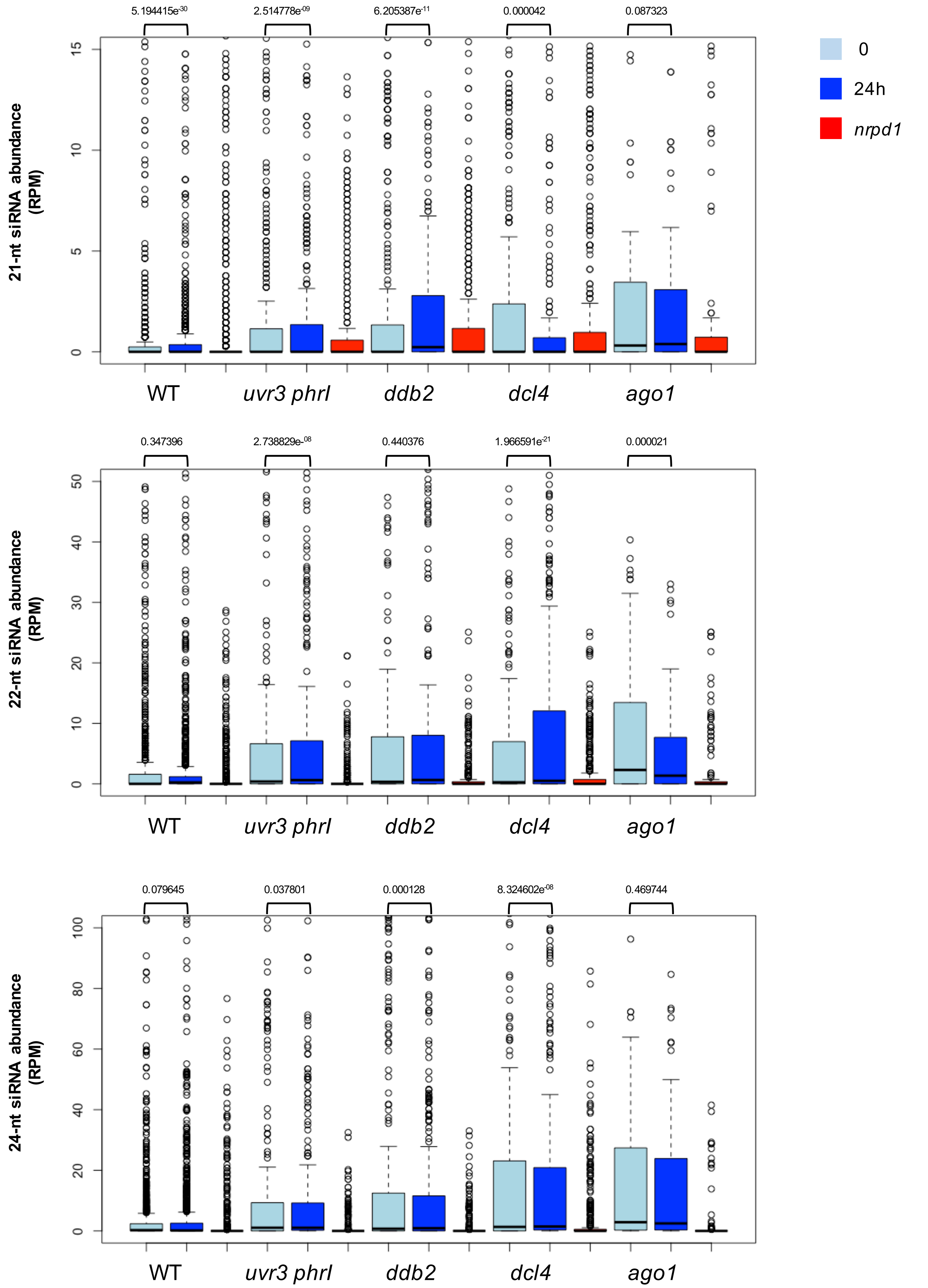


**Supplemental Figure 20**

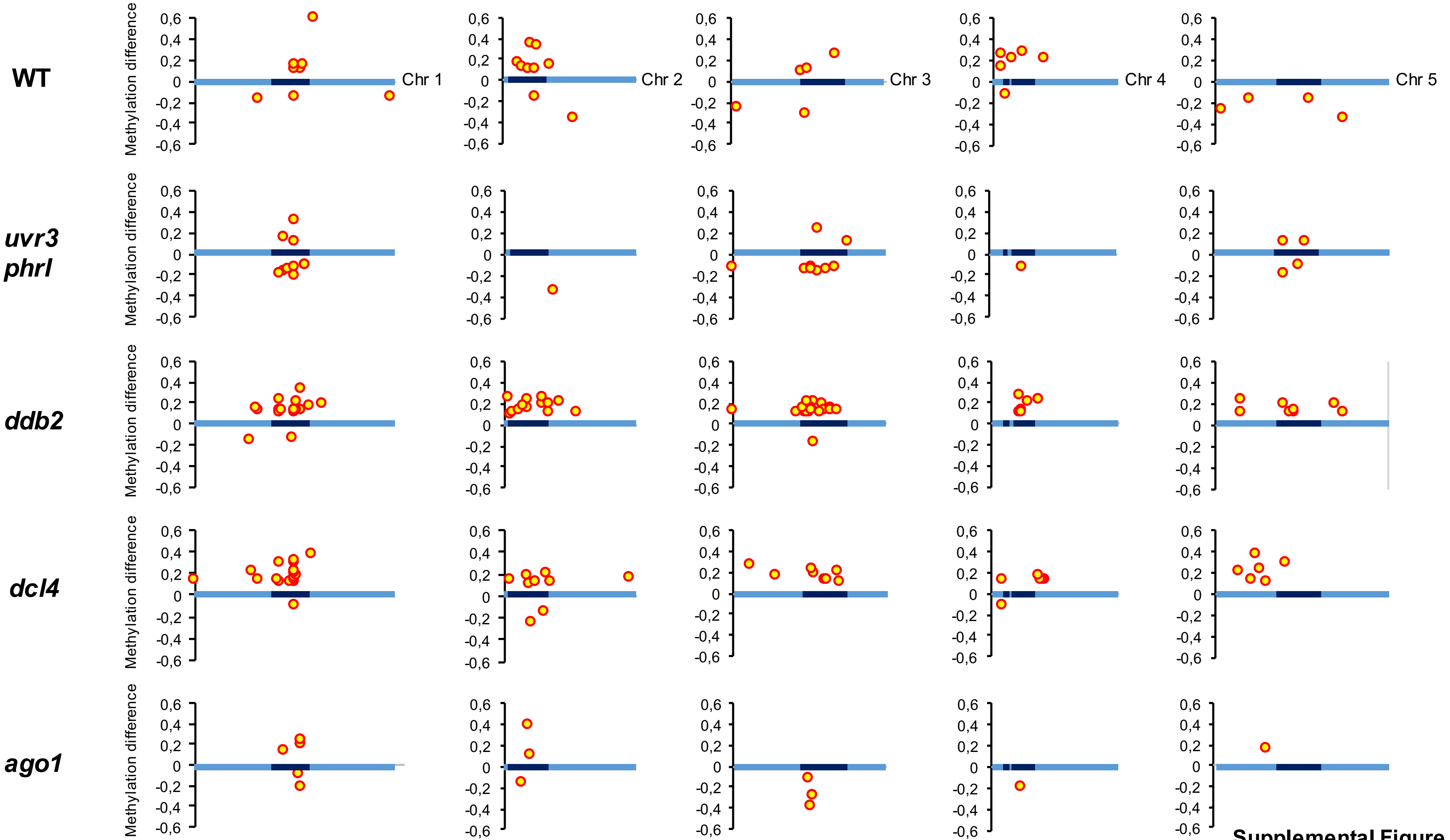


Supplemental Figure 21

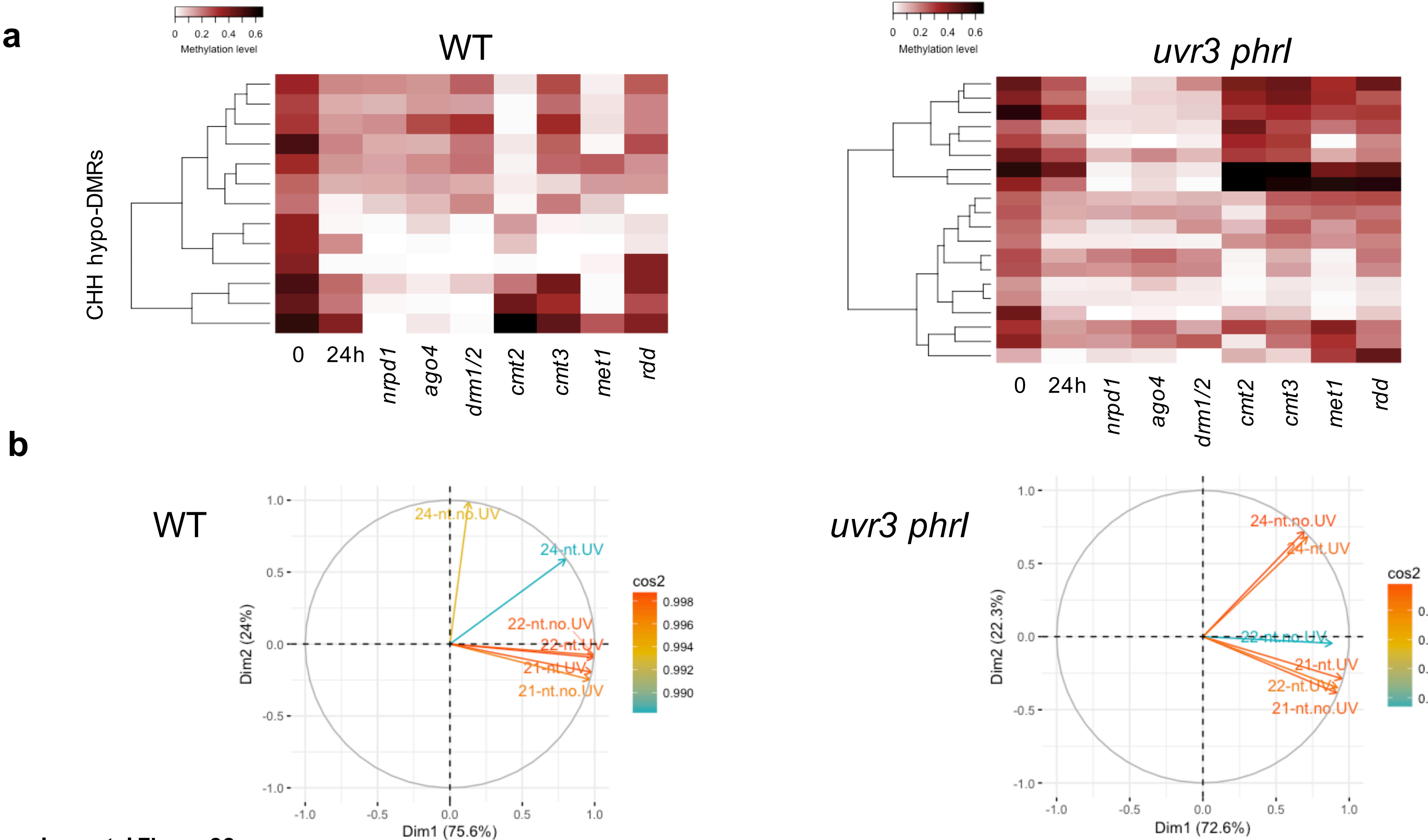


**Supplemental Figure 22**

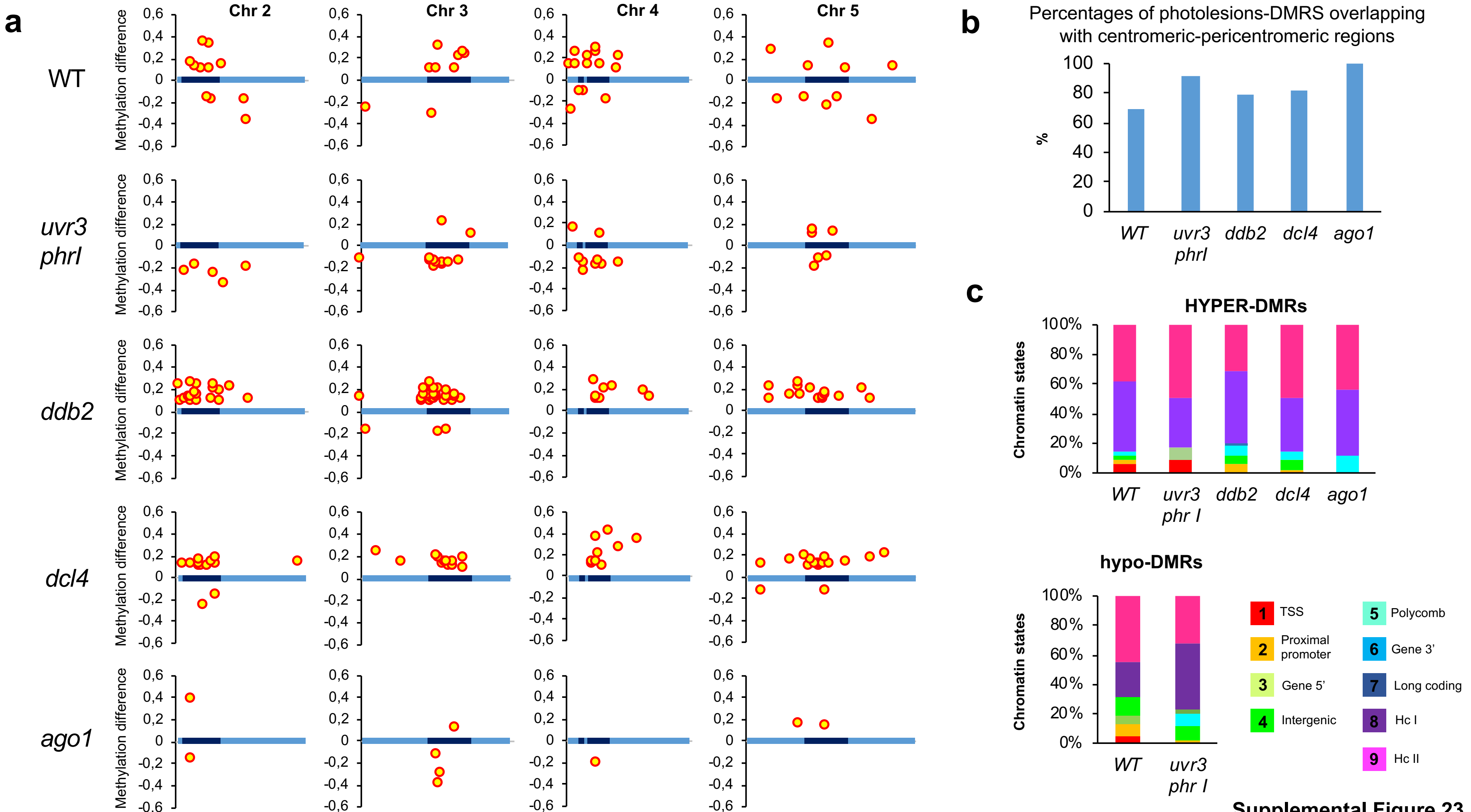


**Supplemental Figure 23**

A

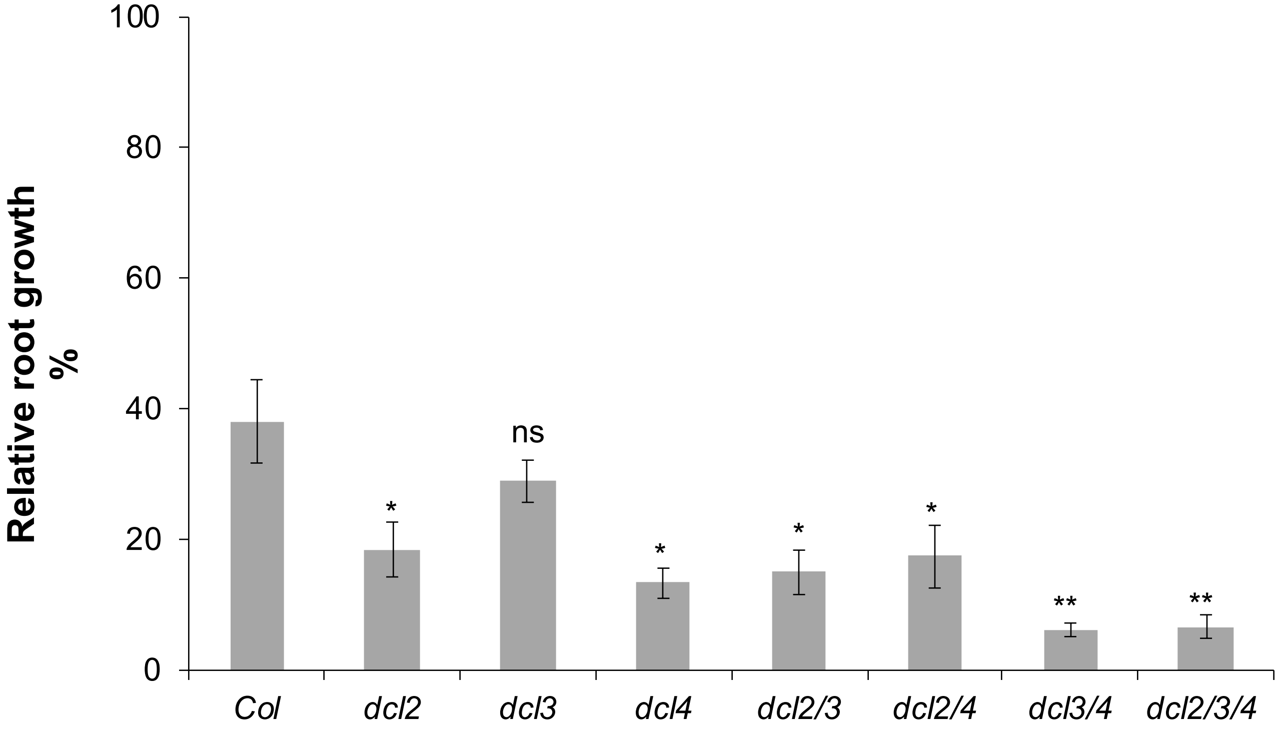


B

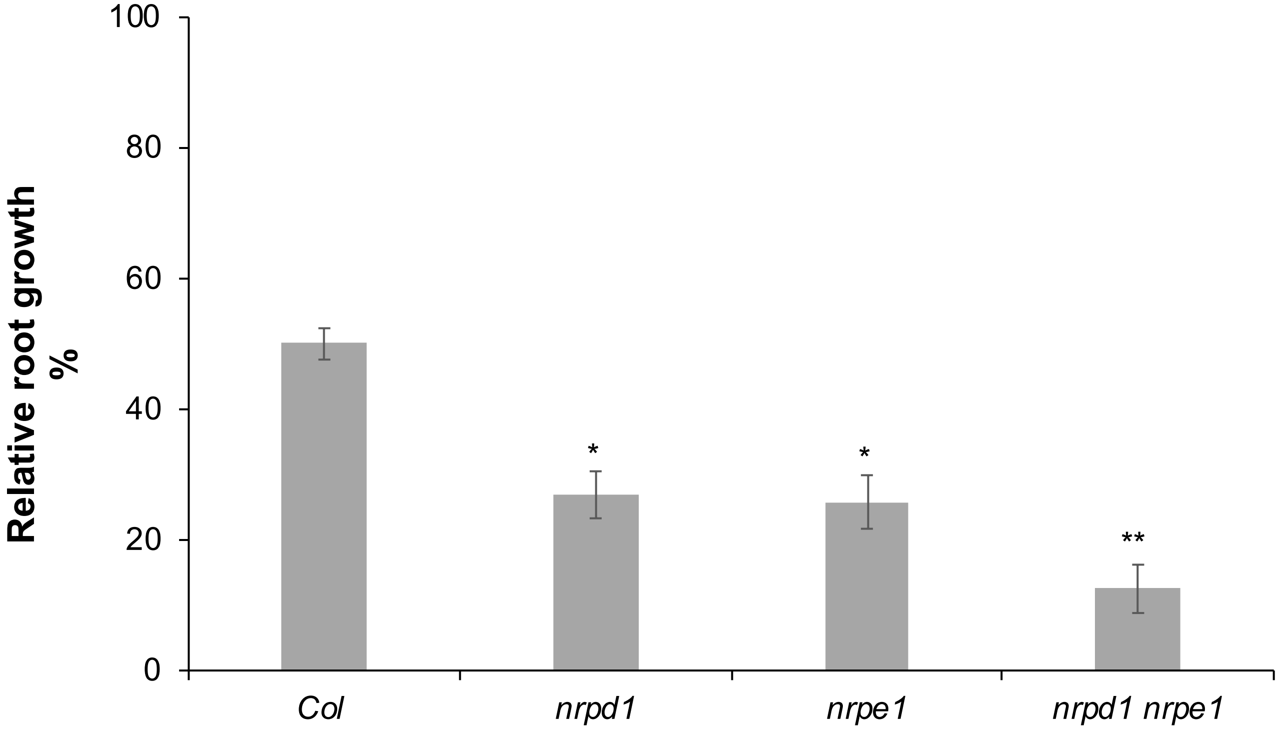


Supplemental Figure 24

**Supplemental Table 1 : Statistics of di-pyrimidines frequencies**

Di-pyrimidine frequencies for each DNA strand (+ and -) of damaged intergenic, TE and Protein Coding Genes (PCGs) regions were calculated and compared to di-pyrimidine frequencies of the Arabidopsis thaliana genome (TAIR10). Exact p values calculated according Mann Whitney test.

red: higher compared to the Arabidopsis genome

blue: lower compared to the Arabidopsis genome

black : non significant

| | **WT** | ***uvr3 phrI*** | ***ddb2*** | ***dcl4*** | ***ago1*** |
|---|---|---|---|---|---|
| **CC+ inter** | 0.000002 | $4.108066e^{-22}$ | $1.043065e^{-11}$ | $1.047994e^{-12}$ | 0.073404 |
| **CC- inter** | 0.001245 | $1.034353e^{-22}$ | $1.181482e^{-07}$ | $2.386710e^{-10}$ | 0.726298 |
| **CT+ inter** | 0.002855 | 0.002043 | 0.430421 | 0.848763 | 0.773534 |
| **CT- inter** | 0.000021 | 0.005344 | 0.046338 | 0.375911 | 0.657247 |
| **TC+ inter** | 0.193070 | $9.911872e^{-12}$ | 0.000154 | 0.000057 | 0.006827 |
| **TC- inter** | 0.278745 | $4.827580e^{-10}$ | 0.555606 | 0.035694 | 0.315787 |
| **TT+ inter** | $2.437233e^{-07}$ | 0.378616 | $1.148263e^{-08}$ | 0.000384 | 0.000056 |
| **TT- inter** | $8.129112e^{-08}$ | 0.009999 | 0.012661 | 0.000759 | 0.230810 |
| **CC+ TE** | 0.000159 | 0.026875 | 0.678068 | 0.880010 | 0.034961 |
| **CC- TE** | 0.325881 | $1.127627e^{-37}$ | 0.000002 | $3.445125e^{-15}$ | 0.006277 |
| **CT+ TE** | $5.181050e^{-20}$ | 0.131896 | 0.009273 | 0.004097 | 0.001526 |
| **CT- TE** | 0.032106 | $3.569692e^{-10}$ | 0.054896 | $2.674007e^{-07}$ | 0.497429 |
| **TC+ TE** | 0.461676 | $4.164820e^{-20}$ | 0.000007 | $1.740909e^{-12}$ | 0.000724 |
| **TC- TE** | 0.140003 | $3.390645e^{-19}$ | 0.000216 | $8.813322e^{-15}$ | 0.080409 |
| **TT+ TE** | $5.371071e^{-36}$ | $1.553131e^{-11}$ | $2.518669e^{-13}$ | $4.612514e^{-14}$ | $1.599210e^{-13}$ |
| **TT- TE** | $1.429321e^{-18}$ | 0.000010 | 0.000022 | 0.110040 | 0.340116 |
| **CC+ PCG** | $1.598449e^{-165}$ | $2.764503e^{-90}$ | $1.025532e^{-64}$ | $7.759190e^{-88}$ | $1.410070e^{-14}$ |
| **CC- PCG** | $5.584896e^{-159}$ | $6.751455e^{-68}$ | $9.031742e^{-51}$ | $1.114985e^{-87}$ | $2.692716e^{-07}$ |
| **CT+ PCG** | $2.425401e^{-07}$ | $1.300389e^{-29}$ | $4.099648e^{-10}$ | $2.583149e^{-19}$ | 0.000010 |
| **CT- PCG** | $2.805784e^{-12}$ | 0.000316 | 0.000008 | 0.014012 | 0.005046 |
| **TC+ PCG** | $3.355920e^{-34}$ | $9.442238e^{-50}$ | $1.392673e^{-24}$ | $1.793070e^{-30}$ | $1.182933e^{-11}$ |
| **TC- PCG** | $1.466859e^{-35}$ | $3.508585e^{-08}$ | $3.041257e^{-11}$ | $3.126330e^{-10}$ | 0.000151 |
| **TT+ PCG** | $7.635052e^{-15}$ | 0.224611 | 0.096155 | 0.000001 | $2.044536e^{-08}$ |
| **tt- PCG** | $1.162764e^{-18}$ | $3.253269e^{-09}$ | 0.728806 | $2.391625e^{-14}$ | 0.003074 |

**Supplemental Table 2:** Statistics of the high throughput sequencing experiments

Whole Genome Bisulfite Sequencing (WGBS)

| Sample | Number of reads | Clean reads | Mapped reads |
|---|---|---|---|
| WT 0 | 24 799 299 | 23 697 656 | 5 515 940 (23.3%) |
| *ddb2* 0 | 39 089 922 | 37 459 779 | 27 508 313 (73.4%) |
| *uvr3 phrI* 0 | 55 452 923 | 52 683 646 | 47 110 980 (89.4%) |
| *dcl4* 0 | 54 254 262 | 51 825 418 | 47 394 165 (91.4%) |
| *ago1* 0 | 50 337 222 | 48 156 436 | 39 258 943 (81.5%) |
| WT 24h | 52 347 786 | 49 702 705 | 45 580 743 (91.7%) |
| *ddb2* 24h | 53 297 047 | 51 321 223 | 45 627 472 (88.9%) |
| *uvr3 phrI* 24h | 46 756 341 | 44 973 274 | 40 559 204 (90.2%) |
| *dcl4* 24h | 52 542 646 | 50 125 786 | 46 528 825 (92.8%) |
| *ago1* 24h | 52 747 953 | 50 711 267 | 46 211 254 (91.1%) |

Immuno-Precipitation Of UV-damaged DNA (IPOUD)

| Sample | Number of reads | Clean reads | Mapped reads |
|---|---|---|---|
| input | 9 395 384 | 8 384 750 | 4 682 424 (55.8%) |
| WT 6,4 PP | 11 293 498 | 10 466 543 | 257 258 (2.5%) |
| WT CPD | 18 038 164 | 17 080 641 | 406 355 (2.4%) |
| *ddb2* 6,4 PP | 15 931 588 | 14 955 429 | 446 234 (3%) |
| *ddb2* CPD | 18 279 550 | 17 380 256 | 280 916 (1.6%) |
| *uvr3 phrI* 6,4 PP | 16 153 008 | 15 286 810 | 499 430 (3.3%) |
| *uvr3 phrI* CPD | 13 978 557 | 13 003 628 | 427 924 (3.3%) |
| *dcl4* 6,4 PP | 14 332 390 | 13 513 948 | 545 923 (4.0%) |
| *dcl4* CPD | 14 974 110 | 14 148 505 | 290 613 (2.0%) |
| *ago1* 6,4 PP | 13 300 629 | 12 388 736 | 193 769 (1.6%) |
| *ago1* CPD | 18 128 703 | 17 105 325 | 137 551 (0.8%) |

Small RNA-seq

| Sample | Number of reads | 15- to 41-nt reads | Mapped reads |
|---|---|---|---|
| WT 0 | 19 677 898 | 14 992 007 | 12 695 174 (84.6%) |
| *ddb2* 0 | 21 989 130 | 17 235 777 | 14 532 863 (84.3%) |
| *uvr3 phrI* 0 | 19 640 581 | 15 254 723 | 12 585 739 (82.5%) |
| *dcl4* 0 | 22 671 494 | 16 281 510 | 13 022 642 (79.9%) |
| *ago1* 0 | 19 688 909 | 16 092 269 | 11 775 752 (73.1%) |
| WT 24h | 27 525 582 | 24 196 187 | 20 193 020 (83.4%) |
| *ddb2* 24h | 22 315 226 | 17 644 910 | 14 194 205 (80.4%) |
| *uvr3 phrI* 24h | 21 924 679 | 18 850 437 | 15 544 956 (82.4%) |
| *dcl4* 24h | 23 111 040 | 18 963 195 | 15 554 742 (82.0%) |
| *ago1* 24h | 18 586 123 | 12 437 522 | 10 194 498 (81.9%) |

**Supplemental Table 3 :** Primers sequences used for qPCR

| Name | Sequence |
|---|---|
| MET1-fw | CGTGGTTCCAAAAGGAGATAAG |
| MET1-rv | TGTTTGGGAGACAAAAAGGAAT |
| DRM2-fw | TCGTTTCAGGTTGATACTGTGG |
| DRM2-rv | TGGTTAGTTTGCTCCCAAAAGT |
| CMT3-fw | AGGGTCAAACTTGAATCTGGAA |
| CMT3-rv | GATAAAACCCGATTTTGCTCTG |
| GAPDH -fw | TTGGTGACAACAGGTCAAGCA |
| GAPDH-rv | AAACTTGTCGCTCAATGCAAT |
| UbiCRed-fw | ACAAGCCAATTTTTGCTGAGC |
| UbiCRed-rv | ACAACAGTCCGAGTGTCATGGT |
| Hexo-fw | GGCGTTTTCTGATAGCGAAAA |
| Hexo-rv | ATGGATCAGGCATTGGAGCT |
| ROS1-fw | AAGGTCACATGTTGTGAACCAAT |
| ROS1-rv | ATGCTCGTCTGGAAGTTCGTA |
| RTPCR5S1 | GGATGCGATCATACCAG |
| 5SUNIV1 | CGAAAAGGTATCACATGCC |
| 180(all)-F | ACCATCAAAGCCTTGAGAAGCA |
| 180(all)-R | CCGTATGAGTCTTTGTCTTTGTATCTTCT |
| DML3-fw1 | GGACAATTCAGGATTCTTTCAGA |
| DML3-rv1 | CTTCACTAAATGTGATGGAGCATT |
| DML2-fw1 | TCTGTATCCTCGATTTGTAAAGGTT |
| DML2-rv1 | CCTTACACAGACATATCCTTCCTG |
| CMT2-fw | GATCACAGGCCGTTCCATATAA |
| CMT2-rv | ACAGTTTCATCCCACCAAAGA |
| COPIA78F_RT | CCACAAGAGGAACCAACGAA |
| COPIA78R_RT | TTCGATCATGGAAGACCGG |
